# Diurnal.plant.tools: comparative transcriptomic and co-expression analyses of diurnal gene expression of the Archaeplastida kingdom

**DOI:** 10.1101/658559

**Authors:** Jonathan Wei Xiong Ng, Qiao Wen Tan, Camilla Ferrari, Marek Mutwil

## Abstract

Almost all organisms coordinate some aspects of their biology through the diurnal cycle. Photosynthetic organisms, and plants especially, have established complex programs that coordinate physiological, metabolic and developmental processes with the changing light. The diurnal regulation of the underlying transcriptional processes is observed when groups of functionally related genes (gene modules) are expressed at a specific time of the day. However, studying the diurnal regulation of these gene modules in the plant kingdom was hampered by the large amount of data required for the analyses. To meet this need, we used gene expression data from 17 diurnal studies spanning the whole Archaeplastida kingdom (Plantae kingdom in the broad sense) to make an online diurnal database. We have equipped the database with tools that allow user-friendly cross-species comparisons of gene expression profiles, entire co-expression networks, co-expressed clusters (involved in specific biological processes), time-specific gene expression, and others. We exemplify how these tools can be used by studying three important biological questions: (i) the evolution of cell division, (ii) the diurnal control of gene modules in algae and (iii) the conservation of diurnally-controlled modules across species. The database is freely available at http://diurnal.plant.tools/.

## INTRODUCTION

Photosynthetic organisms use two primary systems, photoreceptors and the circadian clock to detect light and measure time (and thus the photoperiod) respectively (Serrano-Bueno *et al.*, 2017). Light quality and quantity is detected by photoreceptors, which have evolved across different photosynthetic organisms. A broad range of photoreceptors can elicit different responses via light signal transduction, such as flowering, photomorphogenesis or circadian rhythms. The circadian clock is an internal, self-sustained system based on feedback-loops and other factors that generate a timekeeping network and synchronizes light detection with other biological processes in the plant (Nohales and Kay, 2016). Diurnal rhythms, which also include non-endogenous oscillations, are found in almost all eukaryotes, repeat roughly every 24 hours, and are governed by highly similar gene networks across different species (Serrano-Bueno *et al.*, 2017; Nohales and Kay, 2016).

In photosynthesizing organisms, light is the major signal for diurnal rhythms, as many basic biological processes, such as carbon fixation and energy supply are driven by light signalling. Consequently, many other biological processes, such as cell division, metabolic activity and growth, are regulated by alternating light and dark intervals (Covington *et al.*, 2008; Bell-Pedersen *et al.*, 2005; Zones *et al.*, 2015; de los Reyes *et al.*, 2017). Diurnal regulation of these biological processes is often under strong transcriptional control. For example, in the alga *Ostreococcus tauri*, mRNA levels of genes involved in cell division, the Krebs cycle and protein synthesis are under control of light/dark cycles (Monnier *et al.*, 2010; de los Reyes *et al.*, 2017). On the other hand, gene expression, RNA editing, splicing, and noncoding RNA expression are controlled by diurnal cycles in *Drosophila melanogaster* (Hughes *et al.*, 2012). Recent studies on plants have shown a high level of conservation of diurnal responses even among distantly related species (Ferrari *et al.*, 2019). Hence, diurnal gene expression is a universal and fundamental feature of all living organisms.

Genes with similar expression profiles across, for example, environmental perturbations, developmental stages, and organs tend to be functionally related, and the identification of these co-expressed genes is thus a powerful tool to study gene function (Stuart *et al.*, 2003). These co-expressed genes can be represented as networks, where nodes (or vertices) correspond to genes and edges (or links) connect and indicate genes that have similar expression profiles (Lee *et al.*, 2015). Thus, analysis of gene expression data and co-expression networks can reveal groups of functionally related genes (i.e. gene modules), while their conservation over large phylogenetic distances can reveal core components of shared processes (Ruprecht, Proost, *et al.*, 2017; Ruprecht, Vaid, *et al.*, 2017; Ruprecht *et al.*, 2016).

Despite the pervasiveness of diurnal regulation of biological processes in the plant kingdom, the expression, regulation and conservation of the diurnal programs is not well understood and difficult to study without proper tools. To remedy this, we present diurnal.plant.tools, a database containing diurnal transcriptomes of 17 species of Archaeplastida, which includes eukaryotic algae as well as different phylogenetic groups of terrestrial plants, representing the nine major clades of photosynthetic eukaryotes. The database is preloaded with tools to study the features and conservation of diurnal gene expression in the Archaeplastida kingdom.

## RESULTS

The database contains 17 members of the Archaeplastida kingdom (Figure 1A), and offers a wide selection of tools to query the database. For example, the user can find the genes of interest by using sequence similarity (BLAST), gene IDs (e.g. *At5g67100*) and keywords (e.g. photosynthesis). Genes that work together in a specific biological process or contain a particular domain can be identified by querying the database with GO terms (e.g., GO:0015979) or a Pfam domain (e.g., NAC-dom).

**Figure 1.**
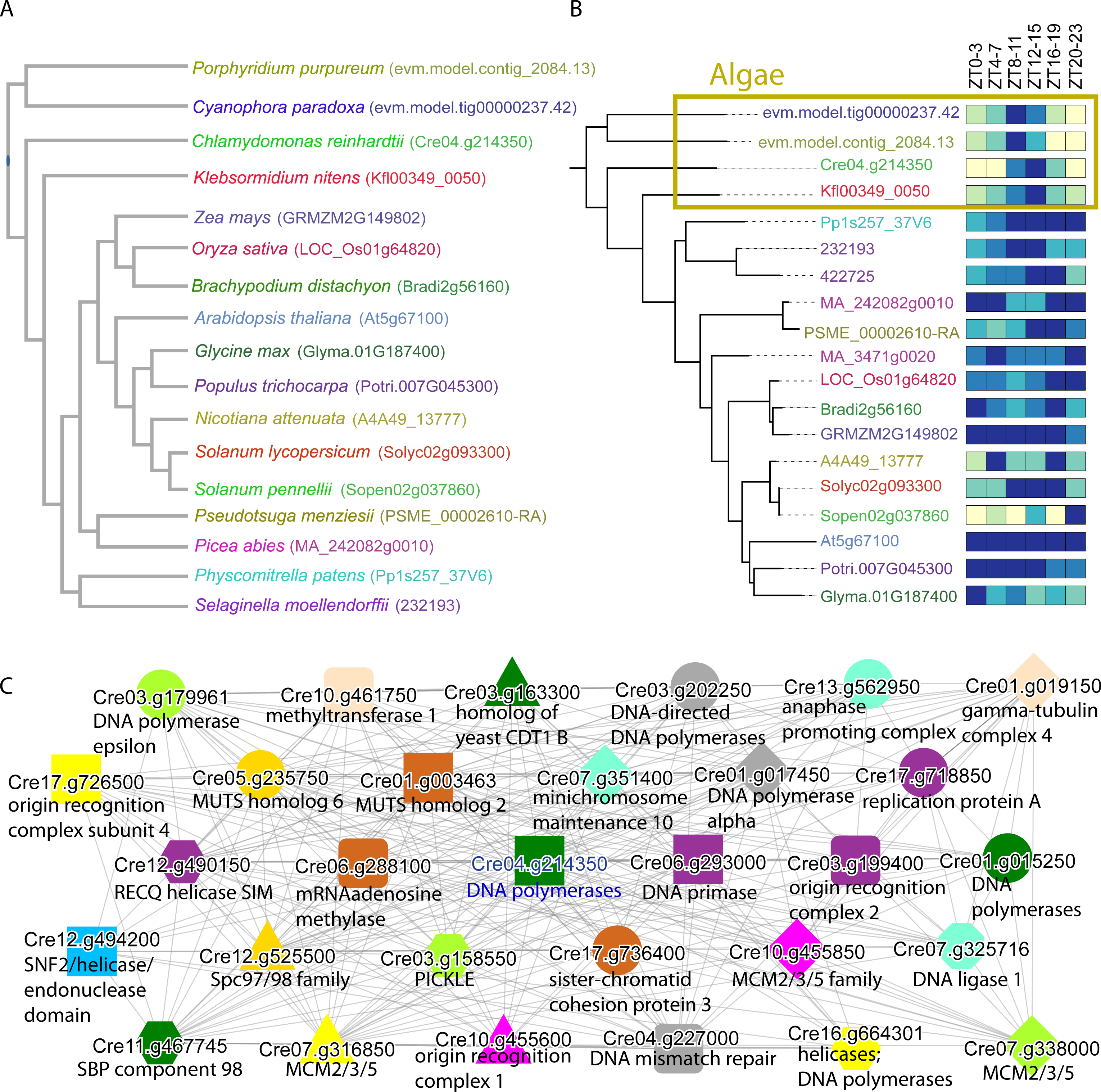
Analysis of cell division in the Archaeplastida kingdom. A) Phylogenetic tree of the species in the database, along with an example of how their gene ID is written. B) Cladogram of the family of DNA polymerases in the different species. The heatmap shows the diurnal expression of the genes, where ZT0-3 indicates expression 0-3 hours after onset of light. Lightly coloured boxes indicate lower gene expression whereas the darker boxes indicate higher gene expression. C) Gene coexpression network showing genes coexpressed with *Cre04.g214350*. The colour and shape of the labels indicate genes belonging to the same orthogroup, and solid lines indicate co-expressed genes.

The database offers multiple comparative genomic and transcriptomic tools that allow the user to view and compare expression profiles within and across species, and to investigate the phylogenetic and diurnal expression relationships of gene families. A full description of the features is found at https://diurnal.sbs.ntu.edu.sg/features. To exemplify some of these features, we provide three typical case studies.

### Cell division is synchronized by light in algae

Algae cultures where cell division is synchronized by light and dark cycles tend to show highly specific temporal expression patterns. For example, in *Chlamydomonas reinhardtii*, transcripts of genes involved in DNA replication and cell division tend to peak specifically at the transition from light to dark (Zones *et al.*, 2015). Our previous study suggested that all algae show this behaviour (Ferrari *et al.*, 2019), indicating that cell division in algae is generally synchronized by light.

To investigate whether diurnal.plant.tools can be used to study the evolution of cell division in the plant kingdom, we examined the expression profile of the *A. thaliana* gene *At5g67100*, which encodes an essential DNA polymerase alpha (Barrero *et al.*, 2007). The expression profile of *At5g67100* (https://diurnal.sbs.ntu.edu.sg/sequence/view/106068) is relatively uniform, which indicates that the gene is not under strong diurnal control (Figure S2), thus suggesting that cell division is not under strong diurnal control in *A. thaliana*.

To investigate the diurnal expression patterns of the DNA polymerases alpha in the plant kingdom, we clicked on the “Gene families: OG_02_0006443” link to arrive at a page containing the sequences, phylogenetic trees, and expression profiles of the genes found in this family (https://diurnal.sbs.ntu.edu.sg/family/view/56123). The page revealed that all species in the database contained at least one copy of the DNA polymerase. By clicking on the *C. reinhardtii* gene *Cre04.g214350* (https://diurnal.sbs.ntu.edu.sg/sequence/view/250705), we observed that the expression of this DNA polymerase peaks after the light phase, confirming that cell division is synchronized by light in *C. reinhardtii*, as previously reported (Figure S2)(Zones *et al.*, 2015).

Diurnal.plant.tools allows the user to view and compare expression profiles of genes across multiple species. The output can be presented in the form of a heatmap (Figure S3), or a phylogenetic tree (Figure 1B). By clicking on the link of the family’s phylogenetic tree (https://diurnal.sbs.ntu.edu.sg/tree/view/11088), we observed that cell division is indeed synchronized by light in algae (yellow box), as these species tend to show a highly specific peak of gene expression around zeitgeber 8-15 (ZT, hours after the light was turned on). Conversely, land plants show a more uniform expression of the cell division genes, suggesting that cell division is not under diurnal control, at least in the sampled tissues.

Since functionally related genes should be expressed with the same spatio-temporal pattern (Zones *et al.*, 2015; Ferrari *et al.*, 2019), we hypothesized that genes important for cell division should be co-expressed. The gene page of *Cre04.g214350* (https://diurnal.sbs.ntu.edu.sg/sequence/view/250705) contains a link to view the co-expression neighborhood network, which represents the gene module that the DNA polymerase is part of (https://diurnal.sbs.ntu.edu.sg/network/graph/81779). In this gene module, many of the genes coexpressed with *Cre04.g214350* are related to DNA replication or cell division (Figure 1C). Genes involved in DNA replication include, among others, DNA polymerase epsilon catalytic subunit (*Cre03.g179961*, light green circle)(Henninger and Pursell, 2014), the origin recognition complex subunit 4 (*Cre17.g726500*, yellow square)(Li *et al.*, 2018) and DNA primase (*Cre06.g293000*, purple square)(Guilliam *et al.*, 2015). Genes involved in cell division include gamma tubulin complex 4 (*Cre01.g019150*, beige diamond)(Janke, 2014), anaphase-promoting complex subunit 6 (*Cre13.g562950*, light blue circle)(Alfieri *et al.*, 2017) and sister-chromatid cohesion protein 3 (*Cre17.g736400*, orange circle)(Losada, 2008).

This neighborhood supports the notion that algae exhibit tight diurnal control of cell division and that co-expression networks based on diurnal gene expression can be used to study diurnally-controlled gene modules.

### Network analysis of diurnal expression of biological processes in *C. reinhardtii*

A recent study in *C. reinhardtii* revealed an intricate diurnal program coordinating the temporal expression of various biological processes in the alga (Zones *et al.*, 2015). To show how diurnal.plant.tools can be used to study temporal regulation of biological processes, we obtained lists of genes involved in various biological processes such as protein synthesis, flagella formation, cell division, respiration and photosynthesis (Table S1)(Zones *et al.*, 2015). These genes were entered into ‘Tools/Create custom network’ (https://diurnal.sbs.ntu.edu.sg/custom_network/), and we selected *C. reinhardtii* as the species and ZT time to color the nodes according to the zeitgeber time (the cytoscape network is available as Supplemental Data 1).

The analysis showed that chloroplast ribosomal subunits are among the first genes to peak (Figure 2, bottom right, red nodes indicate genes peaking at ZT0-3). The co-expression edges between the chloroplast ribosomal small and large subunit genes indicate that these genes are likely controlled by the same transcriptional program. Interestingly, multiple genes representing cytochrome b6f assembly factors and glycolysis/gluconeogenesis-related genes were also co-expressed with the chloroplast ribosomes, suggesting a possible common regulatory pathway controlling these processes. The photosynthesis related genes, such as photosystem I and II subunits (PSI and PSII), light harvesting complexes I and II (LHCI and LHCII), Calvin–Benson–Bassham (CBB) cycle, and cytochrome b6f complex (b6f subunits) reach peak expression at ZT4-11 (predominantly turquoise-yellow nodes).

**Figure 2.**
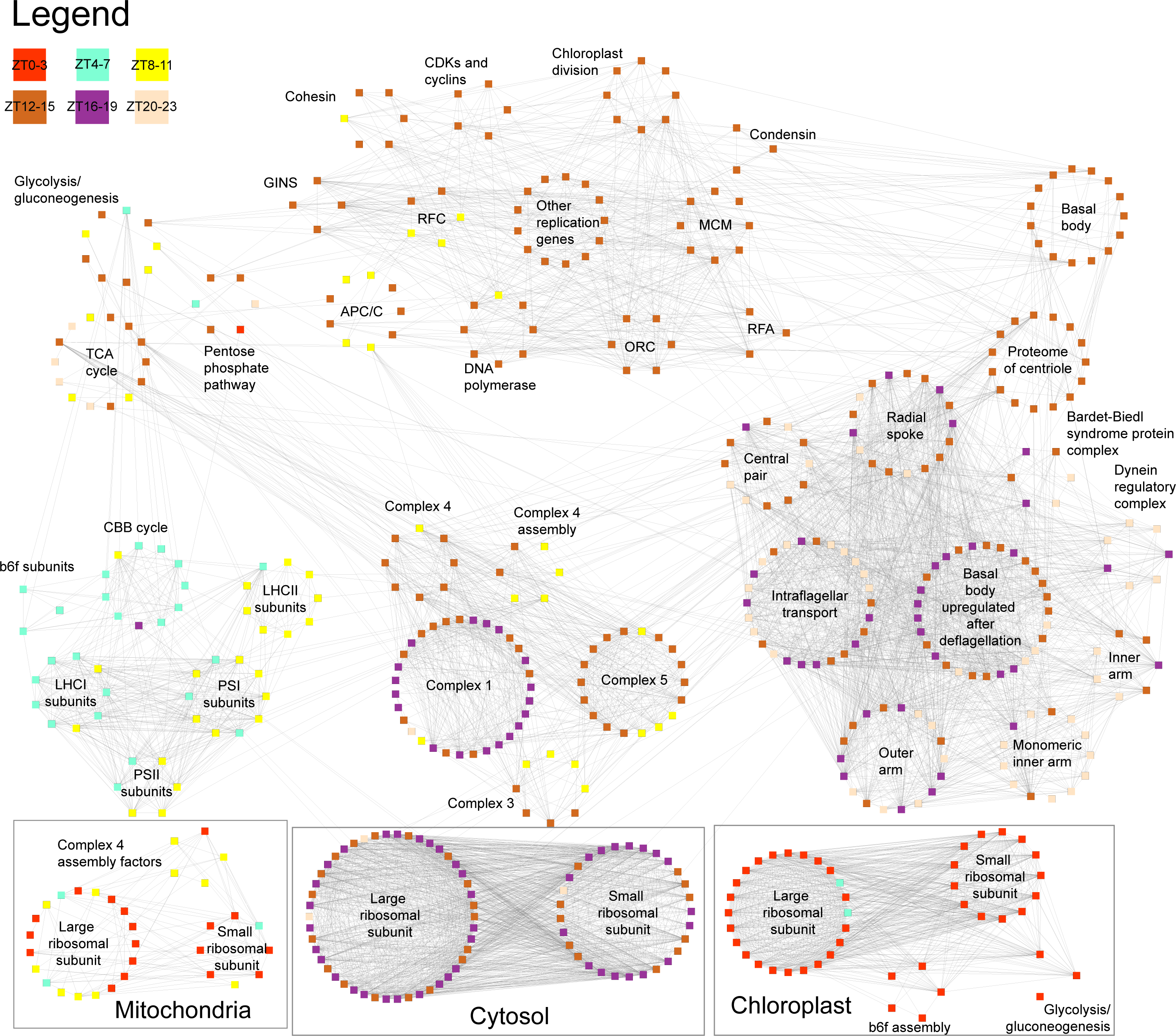
Co-expression network of major biological processes of *C. reinhardtii*. Nodes represent genes and gray edges connect co-expressed genes. The color of the nodes indicates peak expression of a given gene after onset of light (ZT, zeitgeber in hours). For brevity, the gene IDs and disconnected genes are not shown. The Cytoscape 3 network file is available as Supplementary Data 1.

The mitochondrial ribosomal subunits tend to be expressed between ZT0-11 (predominantly red-yellow nodes), indicating that the mitochondrial ribosomal subunits are expressed later than the chloroplast counterparts (Zones *et al.*, 2015). Interestingly, the mitochondrial electron transport complex IV assembly factors are also transcriptionally associated with the ribosomes. The mitochondrial electron transport complexes III, IV and V were co-expressed with each other and their expression peaked during the transition from light to dark (ZT8-15, yellow-brown nodes), with the exception of complex I (ZT12-19), suggesting that complex I is assembled later. The expression of glycolysis/gluconeogenesis, tricarboxylic acid cycle (TCA cycle) and pentose phosphate pathway showed similar expression patterns to the mitochondrial and photosynthesis complexes.

The various cell division components, such as DNA polymerases, cohesins (sister chromatid separation)(Peters *et al.*, 2008), condensins (chromosome assembly and segregation)(Hirano, 2012), chloroplast division factors, and origin recognition complex (ORC), showed a specific expression in the first three hours after onset of darkness (ZT12-15, brown squares). The extracellular components of the flagella are strongly co-expressed and tend to peak after the cell division genes (ZT16-23, purple-beige nodes). This is expected, as the flagella needs to be rebuild after mitosis (Zones *et al.*, 2015). Since the basal bodies act as centrioles during mitosis (Plotnikova *et al.*, 2009; Parker *et al.*, 2010), the basal body proteins and proteome of centrioles peaked at the same time (ZT12-15) and were co-expressed with the cell division genes.

Taken together, the custom network tool can be used to identify temporal expression and co-expression of multiple biological processes, and thus allows the proposal of a chronological order of events for further experimental examination.

### Cross-species comparison of functional gene modules

Since functionally related genes tend to be connected in the co-expression networks, the groups of densely connected genes will form clusters that are enriched for specific functions, gene families and protein domains. Diurnal.plant.tools allows the users to identify clusters that are enriched for GO terms, thus enabling identification of genes relevant for a specific biological process of interest.

To exemplify how functionally enriched clusters are identified, we entered the GO term for photosynthesis (GO:0015979), which redirected us to the corresponding page (https://diurnal.sbs.ntu.edu.sg/go/view/10041). There are in total 5713 genes (sequences) annotated with this GO term and multiple clusters significantly enriched (adjusted p-value < 0.05) for this term across all species in the database, except maize. Each of these clusters thus represent a source for genes related to some aspect of photosynthesis. Clicking on any of the clusters will redirect the user to a page dedicated to the cluster, which contains an average expression profile of the genes found in the cluster, enriched GO terms, a list of similar clusters, the identity of genes in the cluster as well as gene families and protein domains found in the cluster (e.g. *C. reinhardtii* cluster 13 can be accessed at https://diurnal.sbs.ntu.edu.sg/cluster/view/2337).

We extracted one significantly enriched photosynthesis cluster per species and exemplify average expression profiles of these clusters (the cluster with the lowest adjusted p-value was selected, Figure 3A). Interestingly, the expression profiles of these clusters indicated unique expression profiles, with algae showing stronger expression of photosynthesis genes in the light phase (*C. paradoxa, P. purpureum, C. reinhardtii*), with the exception of *K. nitens* (Figure 3A). Conversely, clusters from land plants showed more uniform expression profiles (*A. thaliana, S. moellendorfii, P. menziesii*) or higher expression in light (*P. abies, P. patens, S. lycopersicum, S. pennelii, B. distachyon, G. max*).

**Figure 3.**
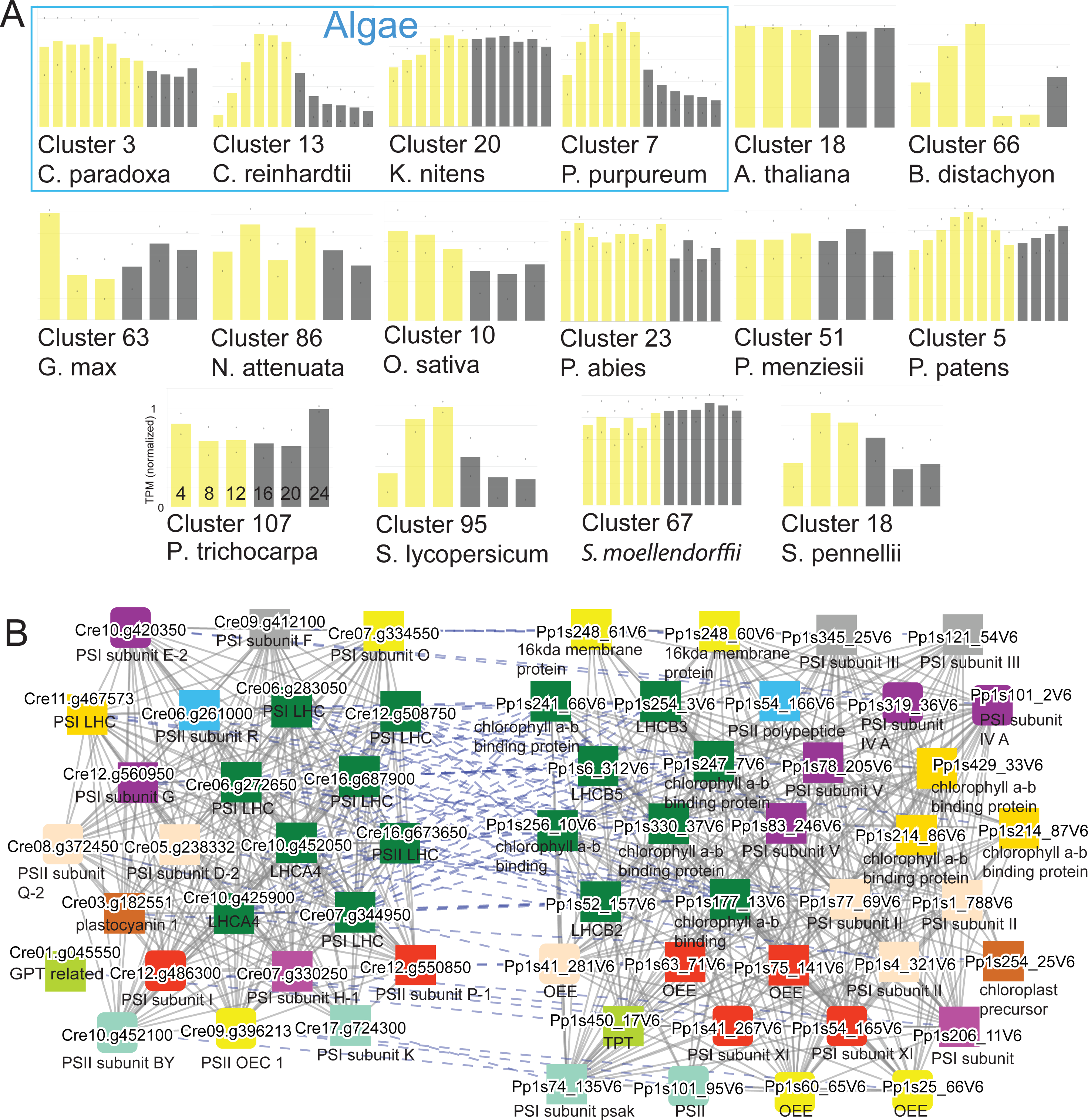
Analysis of photosynthetic clusters. A) Comparative gene expression profiles of gene clusters enriched with the GO term GO:0015979 (photosynthesis) for the 17 species. Yellow bars indicate the light phase, whereas grey bars indicate the dark phase. The bars are ordered according to the zeitgeber time, as exemplified by *P. trichocarpa*. The blue box indicates the algae, while the remaining species are land plants. B) Comparison of the photosynthesis clusters from *C. reinhardtii* (left) with *P. patens* (right). Coexpressed genes are connected with solid gray lines while orthologs found in both neighbourhoods are connected with dashed blue lines. Coloured boxes indicate genes belonging to the same orthogroup.

Next, we compared the photosynthesis gene clusters from *C. reinhardtii* and *P. patens* (Figure 3B) by selecting the *P. patens* Cluster 5 from *C. reinhardtii* cluster 13 page (https://diurnal.sbs.ntu.edu.sg/graph_comparison/cluster/2337/3036/2). Despite the differences in the expression profiles of the clusters of the two species (Figure 3A), and the large evolutionary distance between the species, the two clusters contain similar gene families (Figure 3B). These clusters include genes from photosystems I (PSI) and II (PSII). The PSI genes include PSI subunit E2 (*Cre10.g420350*, purple square)(Ihnatowicz *et al.*, 2007), PSI subunit F (*Cre09.g412100*, grey square)(Nelson and Yocum, 2006) and PSI subunit III (*Pp1s121_54V6*, grey square)(Lotan *et al.*, 1993). The PSII genes include PSII subunit R (*Cre06.g261000*, blue square)(Roose *et al.*, 2016), light-harvesting complex II protein lhcb5 (*Pp1s6_312V6*, green square)(Nelson and Yocum, 2006), and oxygen-evolving enhancer protein 1 (*Pp1s60_65V6*, yellow square)(Heide *et al.*, 2004). To summarize, we have shown through the examples that the cluster enrichment and cluster similarity tools allow users to identify functionally related genes and similar gene modules respectively, with ease.

## CONCLUSIONS

To remedy the lack of resources to study diurnal gene expression in the plant kingdom, we constructed diurnal.plant.tools. This database provides user-friendly tools in form of a modern web-platform that allows users to mine diurnal gene expression data of 17 members of the Archaeplastida kingdom. At the entry level, the users can investigate the diurnal expression of their gene or biological process of interest. For more complex tasks, the plethora of query and analytical tools provide sophisticated methods to generate new hypotheses about the function, evolution and conservation of diurnal gene expression in plants.

## MATERIALS AND METHODS

### Retrieval, annotation and identification of gene families of coding sequences

Coding sequences (CDS) for the 17 species were obtained from different sources as indicated in Table 1. Gene annotations were obtained from their respective online sources, while the CDS of *C. paradoxa, A. thaliana, K. nitens, P. abies, P. purpureum* and *P. menziesii* were annotated using Mercator (Lohse *et al.*, 2014) (http://www.plabipd.de/portal/web/guest/mercator-sequence-annotation).

**Table 1.**
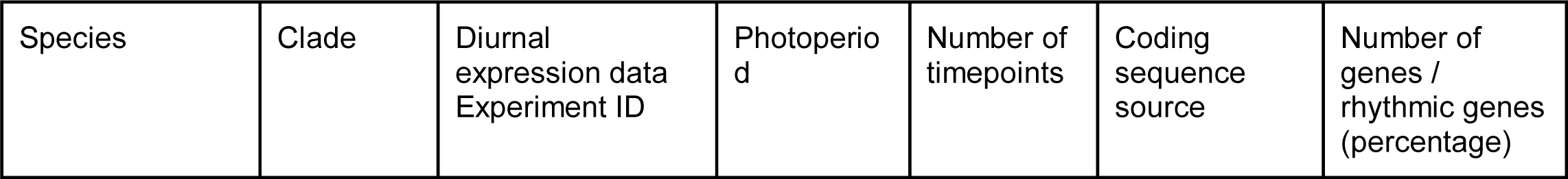
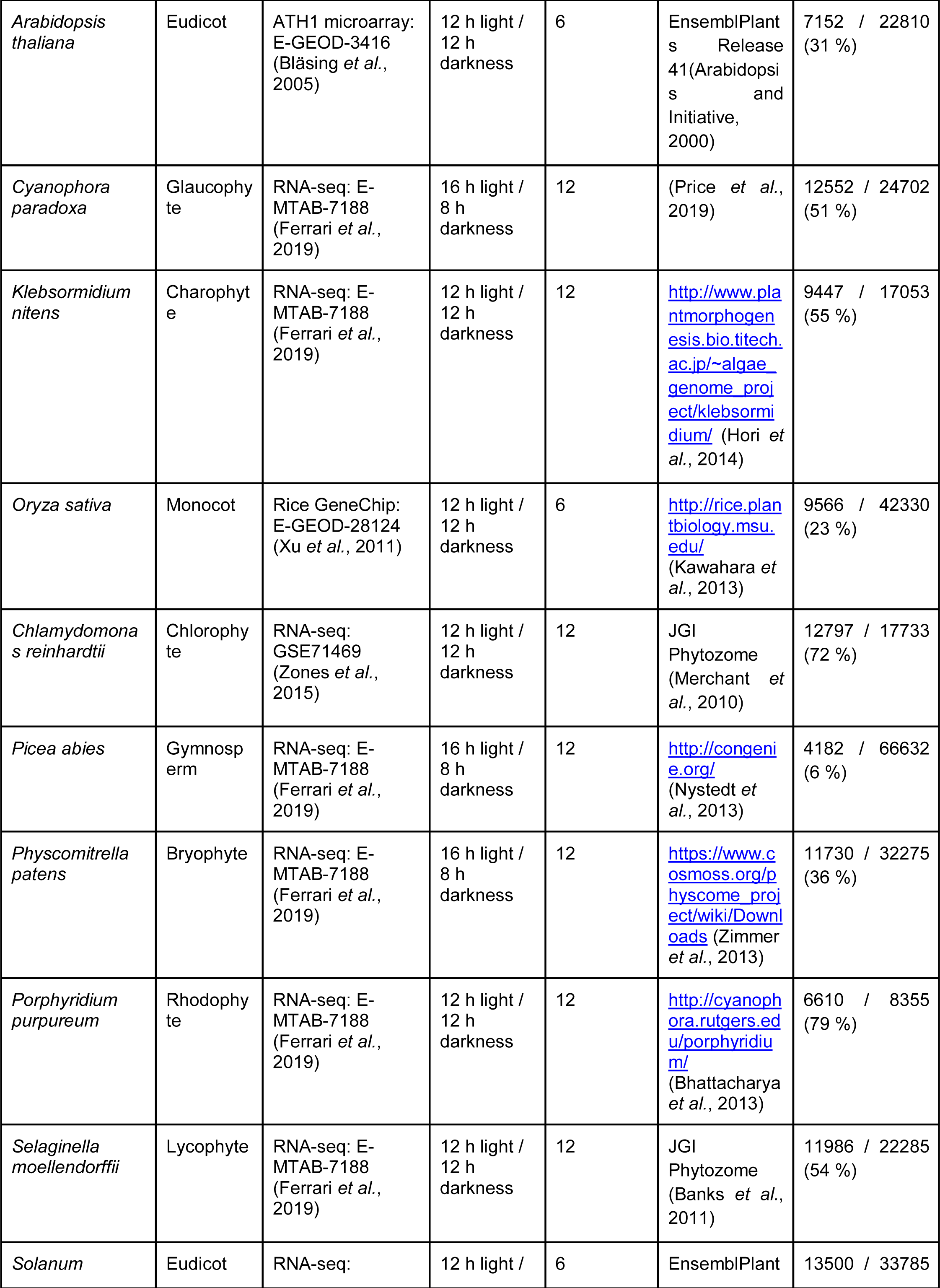
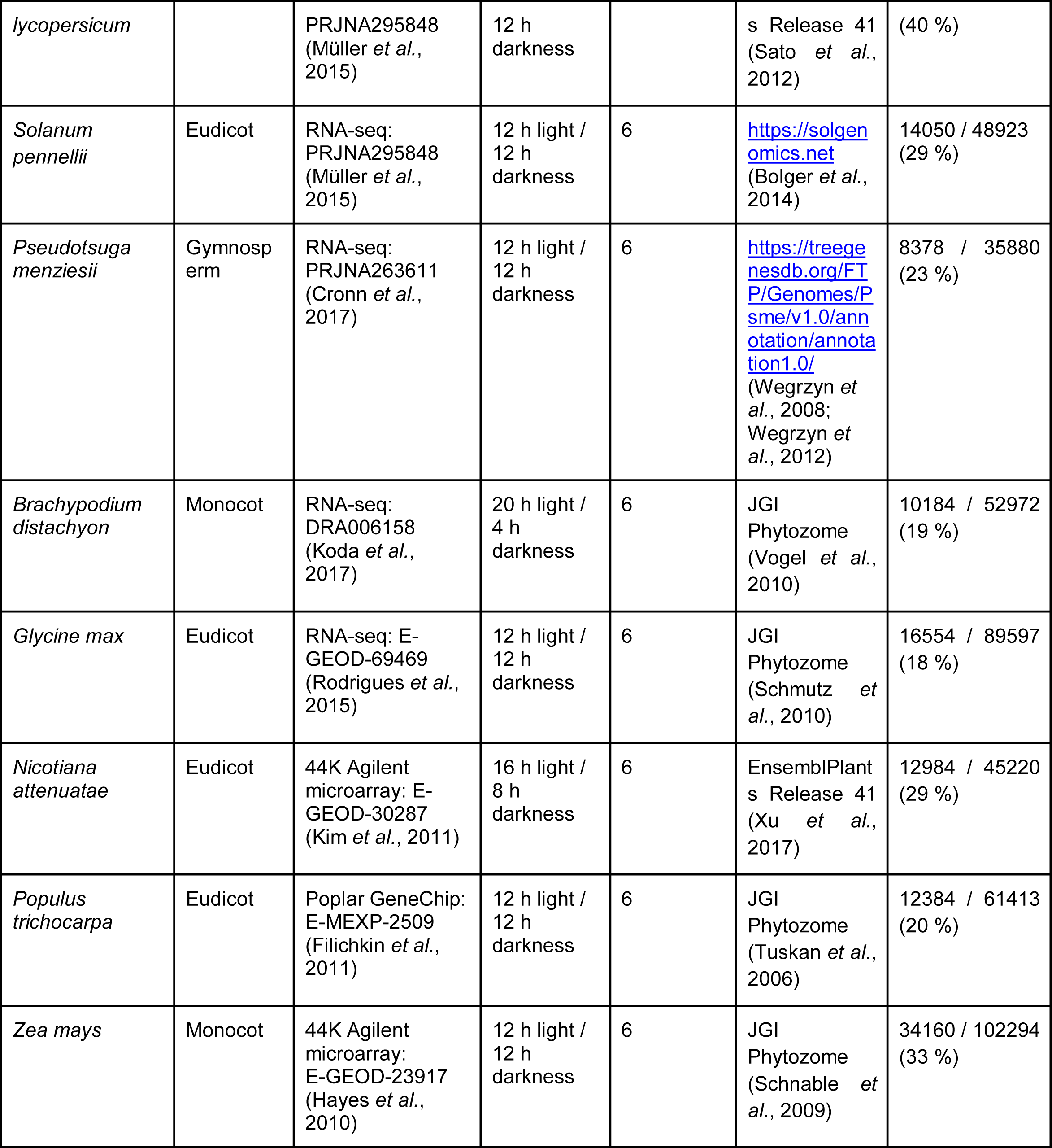
Species and data used to construct diurnal.plant.tools.

Gene Ontology (GO) and protein domain annotations for all organisms except for *S. lycopersicum* were obtained by running InterProScan 5.32-71.0 (Jones *et al.*, 2014) on protein sequences and parsing the Pfam domain and GO term annotations. For *S. lycopersicum*, these annotations were obtained from https://solgenomics.net (Fernandez-Pozo *et al.*, 2015). Gene families were identified by running Orthofinder 1.1.8 (Emms and Kelly, 2015) on the peptide sequences obtained from diurnal.plant.tools.

### Processing gene expression data

Gene expression data capturing diurnal gene expression was collected from sources indicated in Table 1. All RNA-seq data were mapped to their respective CDS and transcripts per million (TPM)-normalized via kallisto(Bray *et al.*, 2016). Genes and probesets from microarray-derived expressions matrices which were not found in the CDS of the respective species were removed from further analysis. Gene expression data for ten species (*A. thaliana, C. paradoxa, K. nitens, O. sativa, C. reinhardtii, P. abies, P. patens, P. purpureum, S. moellendorffii*) were obtained from Ferrari et al. (Ferrari *et al.*, 2019) Principal Component Analysis (PCA) of most of the species revealed that the samples follow an expected circular pattern, where samples representing similar timepoints are adjacent to each other (Figure S1A). The exceptions to this rule were *P. abies* and *P. menziesii*, where the samples did not show a clearly diurnally-dictated grouping of samples.

The JTK_CYCLE (Hughes *et al.*, 2010) algorithm was used to identify genes with significantly (adjusted P-value < 0.05) rhythmic expression patterns from the gene expression matrices. PCA of the gene expression profiles revealed the genes often formed circular patterns, where genes peaking at the same time of the day are adjacent. Similarly to the sample PCA, the exception to the rule were *A. thaliana, P. abies*, and *P. menziesii* (Figure S1B).

The gene expression and gene family data was used to make diurnal.plant.tools database (http://diurnal.plant.tools/), which is based on CoNekT framework (Proost and Mutwil, 2018). Figure 2 was generated by downloading the custom network from diurnal.plant.tools and processing it with Cytoscape 3.7.1 (https://cytoscape.org/). The expression matrices, gene annotation and JTK results for all 17 species can be obtained upon request from the authors.

## DATA AVAILABILITY

The coding and protein sequences are found at https://diurnal.plant.tools.

## ACKNOWLEDGEMENTS

Diurnal.plant.tools is hosted at Nanyang Technological University Singapore and we would like to thank Ryan Chee Kiang Ng for excellent tech support. We would like to thank Dr. Daniela Mutwil-Anderwald for proofreading the manuscript.

## Author Contributions

The database was implemented by J.W.X.N. J.W.X.N., Q.W.T., C.F and M.M. analyzed the data and wrote the manuscript.

## SUPPLEMENTAL TABLES

**Table S1. Gene identifiers and functional categories of *Chlamydomonas* genes used to generate Figure 2.**

## SUPPLEMENTAL DATA

**Data S1. Cytoscape 3 network of the biological processes of *Chlamydomonas reinhardtii.***

## SUPPLEMENTAL FIGURES

**Figure S1.**
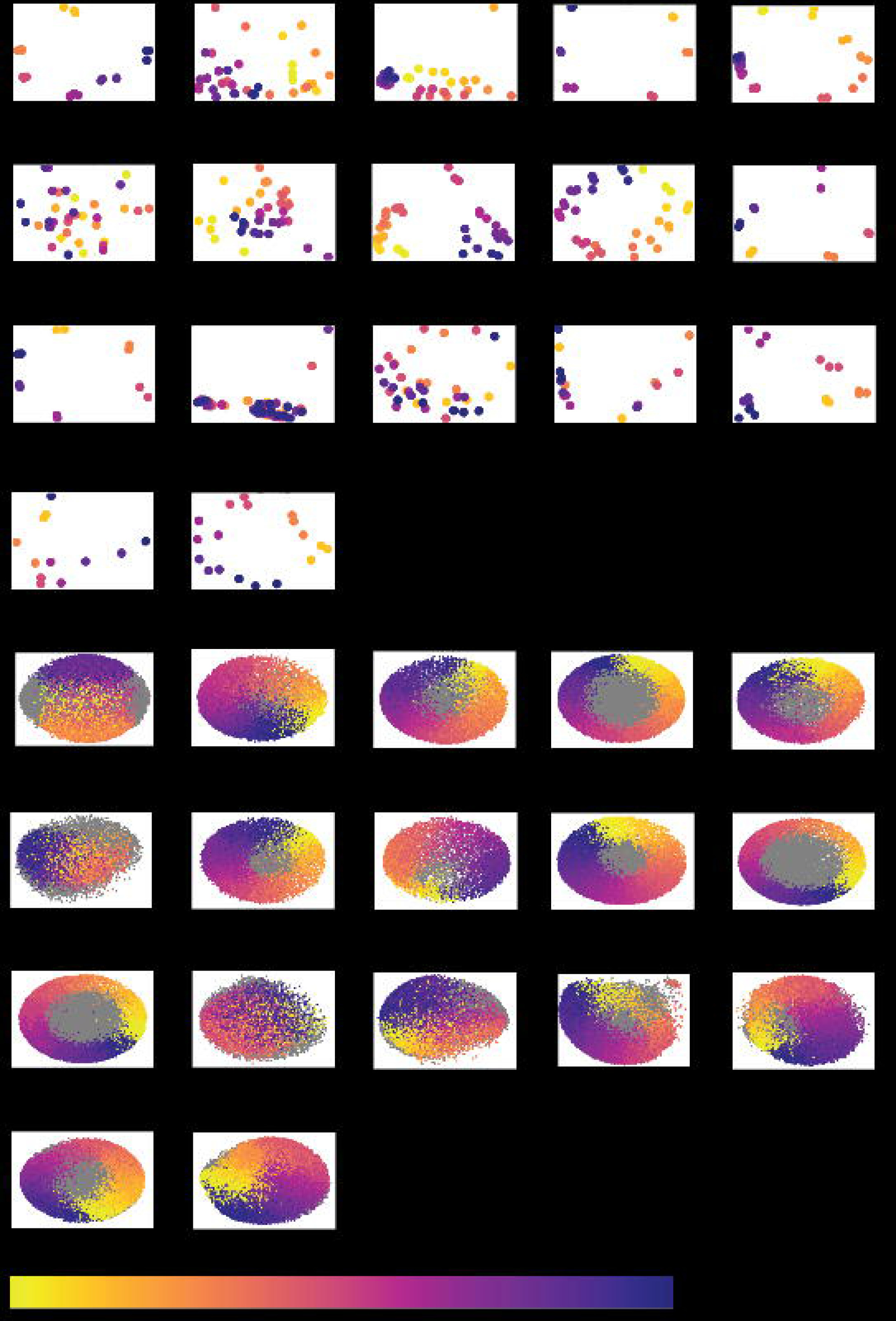
PCA analysis of the samples and genes of the 17 species. A) PCA of the samples. B) PCA of the genes. Each point in the plot corresponds to a sample (A) or a gene (B). The points are color-coded according to the collection time or time of peaking, for samples and genes, respectively, as indicated by the color bar.

**Figure S2.**
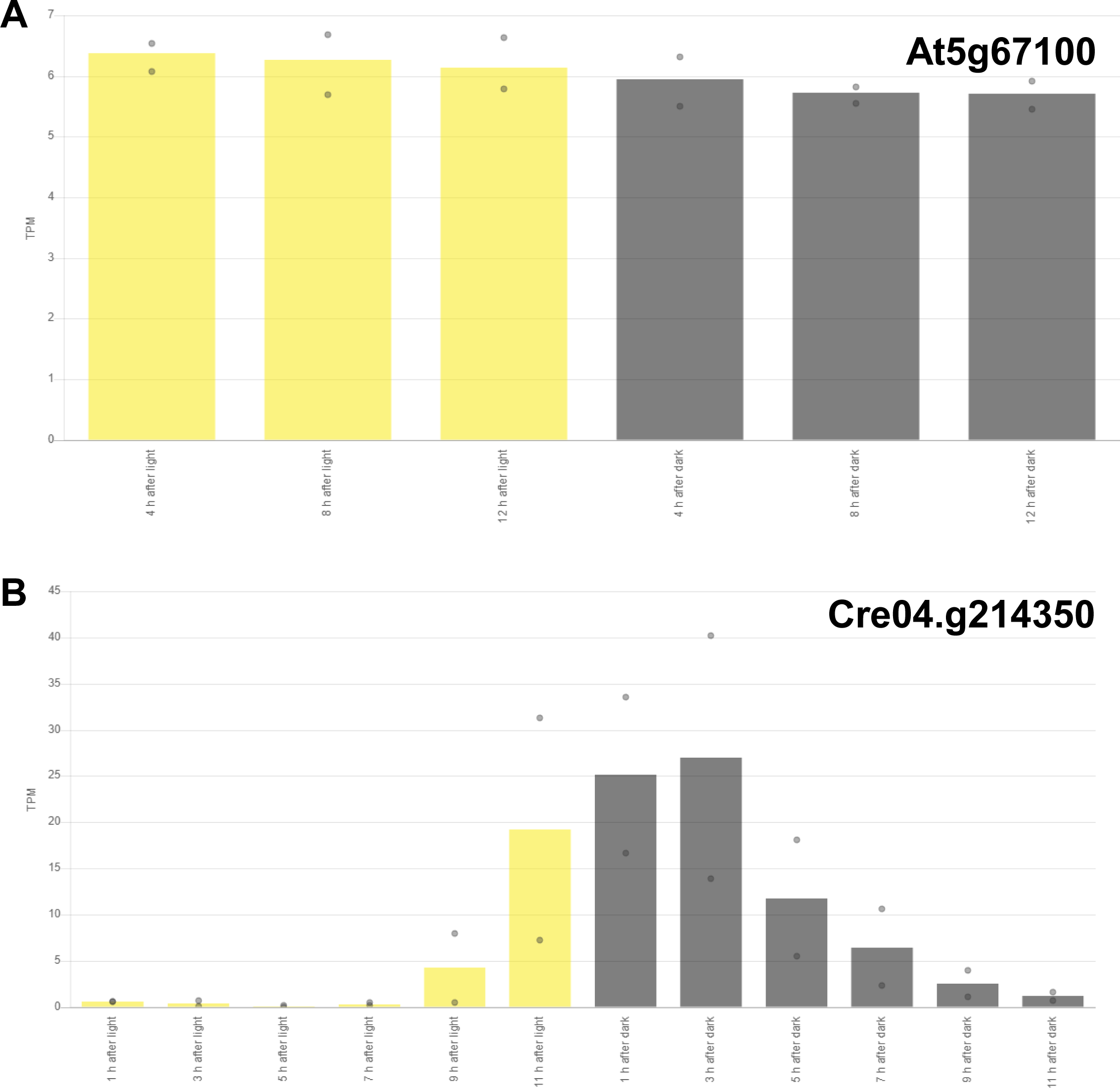
Diurnal expression profiles of DNA polymerase alpha genes from A) *Arabidopsis thaliana* and B) *Chlamydomonas reinhardtii*. The y- and x-axes indicate gene expression levels and the timepoints, respectively. The bars show the average expression levels, while the upper and lower dot indicate the maximum and minimum expression levels, respectively.

**Figure S3.**
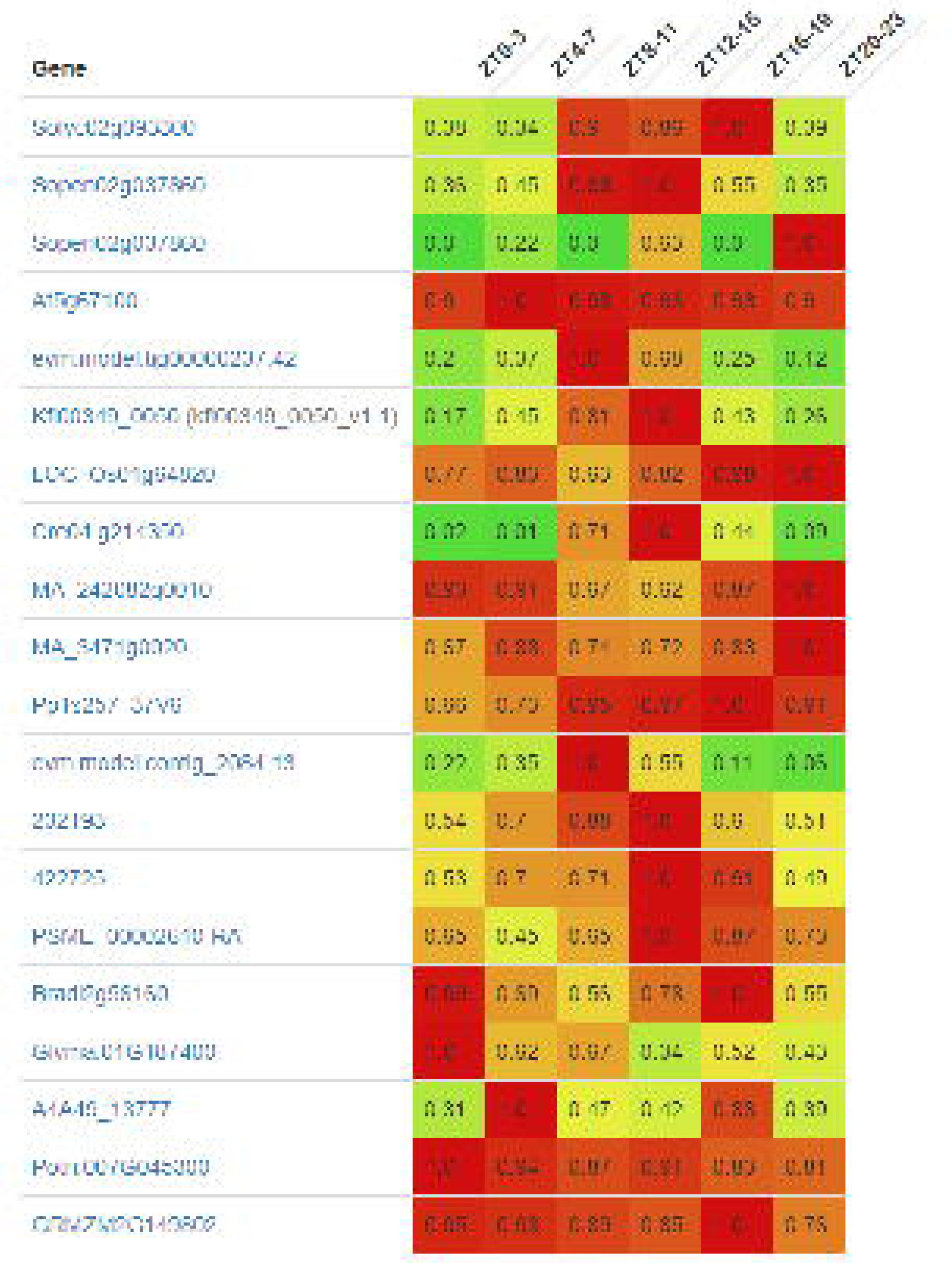
Expression profiles of the genes belonging to OG_02_0006443 gene family. The genes are shown in rows, and the timepoints in columns (as zeitgeber ranges). The coloured cells indicate high (red), low (green) and intermediate (yellow) expression, where each row has been scaled by dividing with the highest expression of the gene.

